# A high-fat diet changes astrocytic metabolism to enhance synaptic plasticity and promote exploratory behavior

**DOI:** 10.1101/2021.11.03.467141

**Authors:** Alexander Popov, Nadezda Brazhe, Anna Fedotova, Alisa Tiaglik, Maxim Bychkov, Kseniia Morozova, Alexey Brazhe, Dmitry Aronov, Ekaterina Lyukmanova, Natalia Lazareva, Li Li, Alexei Verkhratsky, Alexey Semyanov

## Abstract

A high-fat diet (HFD) is generally considered to negatively influence the body, the brain, and cognitive abilities. On the other hand, fat and fatty acids are essential for nourishing and constructing brain tissue. Astrocytes are central for lipolysis and fatty acids metabolism. Here we show that exposure of young mice to one month of HFD elevates lipid content and increases the relative amount of reduced cytochromes in astrocytes but not in neurons. Metabolic changes were paralleled with an enlargement of astrocytic territorial domains due to an increased outgrowth of branches and leaflets. Astrocyte remodeling was associated with an increase in expression of ezrin and with no changes in glial fibrillary acidic protein (GFAP), glutamate transporter-1 (GLT-1), and glutamine synthetase (GS). Such physiological (non-reactive) enlargement of astrocytes in the brain active milieu promoted glutamate clearance and long-term potentiation. These changes translated into improved exploratory behavior. Thus, dietary fat intake is not invariably harmful and might exert beneficial effects depending on the biological context.

**In Brief:** A high-fat diet stimulates the metabolism and growth of astrocytes, which improves glutamate clearance, synaptic plasticity, and exploratory behavior in young mice. Thus, dietary fat arguably is an essential component of the diet for children and young adults, supporting the optimal development of the brain.

**Highlights:**

- Exposure of young mice to a high-fat diet elevated lipid content and increased amount of reduced cytochromes in astrocytes but not in neurons.
- Metabolic changes were paralleled with an enlargement of astrocytic territorial domains due to an increased outgrowth of branches and leaflets.
- Astrocytic enlargement was associated with increased expression of ezrin but not GFAP, hence was not reactive but physiological
- Expansion of astrocytes in the brain active milieu improved glutamate clearance and long-term potentiation.
- The high-fat diet improved exploratory behavior in young mice.

## Introduction

Balanced nourishment is critically essential for brain development, while intake patterns and food composition influence cognitive longevity (Vauzour et al., 2017). Dietary changes impact energetics, morphology, physiology, and life-long adaptive capabilities of the brain. Metabolic homeostasis and metabolic plasticity of brain active milieu (Semyanov and Verkhratsky, 2021) are the fundamental functions of astrocytes. Besides providing the nervous tissue with energy substrates (Belanger et al., 2011) and supplying scavengers of reactive oxygen species (McBean, 2017), astrocytes store glycogen and secrete glycolytic intermediates such as lactate which contribute not only to metabolism but also to intercellular signaling (Cali et al., 2019). Astrocytes are central for the metabolism of lipids, processing the fatty acids, and lipids storage in the form of lipid drops. Astrocytes synthesize sterol lipids, phospholipids, sphingolipids, and cholesterol, all of which are required for producing and maintaining biological membranes (Lee et al., 2021; Pfrieger and Ungerer, 2011). Cholesterol is indispensable for synaptogenesis both in development and adulthood (van Deijk et al., 2017). Astrocytes are the main supplier and neurons primary consumer of cholesterol in the central nervous system (Nieweg et al., 2009). Astrocytic catabolism of fatty acids assists neurons in removing excess of peroxidized fatty acids produced during periods of enhanced neuronal activity(Ioannou et al., 2019). Here we analyzed how high fat intake affects astrocytes and astrocyte-dependent neuroplasticity in young mice.

## Results

### 1. HFD induces metabolic changes in hippocampal astrocytes but not in neurons

Experiments were performed on two groups of mice taken at two months and kept for one month on a standard diet (control) or HFD (Fig. 1a). The weight gain was higher in HFD group (control: 113.5 ± 2.5 %, n = 22; HFD: 128.7 ± 4.0 %, n = 14; p = 0.002, two-sample t-test; Fig. 1b). The extra weight in the HFD group reflected higher body fat percentage (control: 3.0 ± 0.4 %, n = 5; HFD: 8.3 ± 1.0 %, n = 6; p = 0.001, two-sample t-test; Fig. 1c and S1) but not an increase in the weight of viscera. This suggests that HFD animals accumulate more fat in adipose tissue than control mice rather than growing internal organs faster.

**Figure 1.**
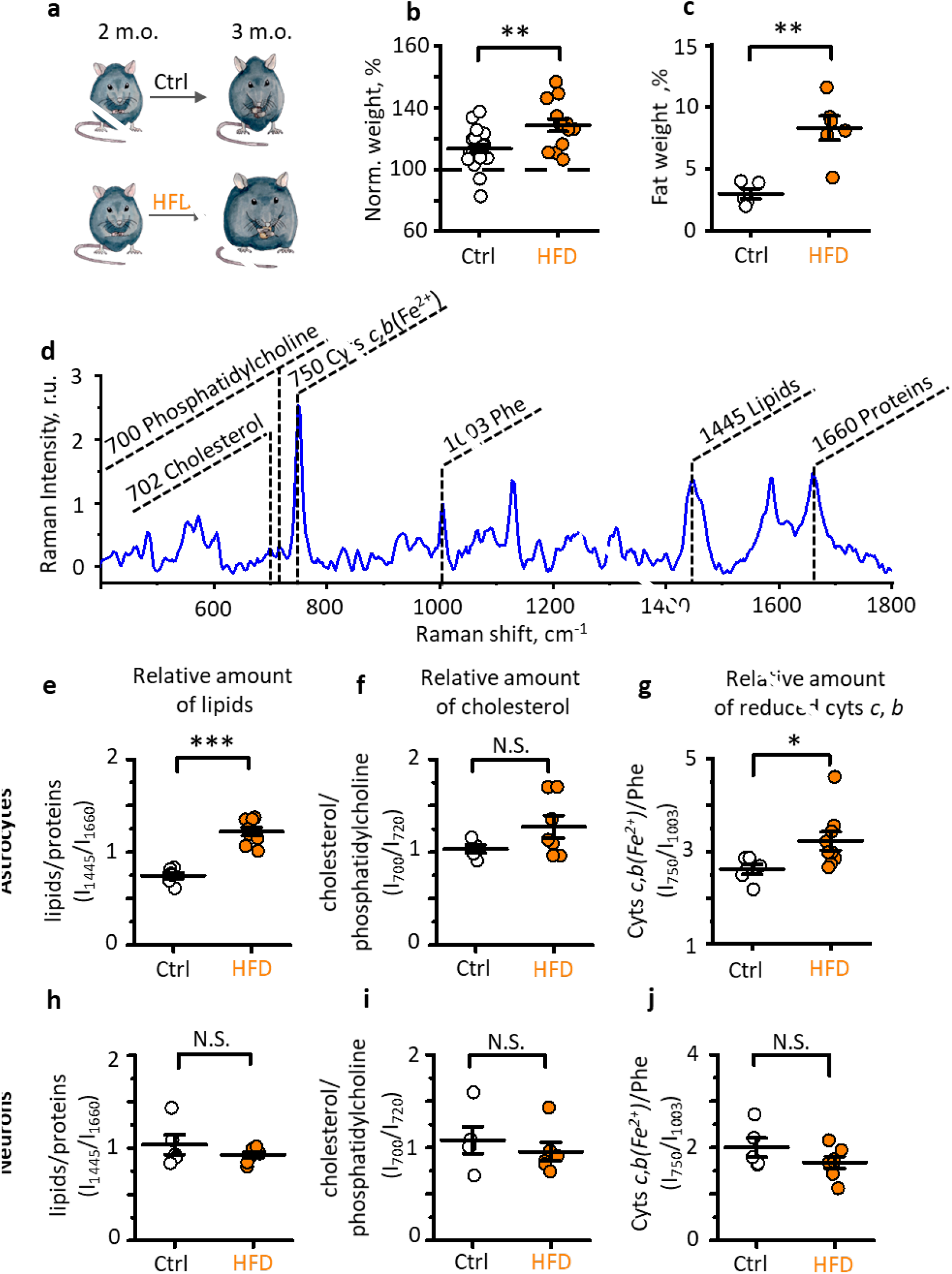
HFD induces metabolic changes in astrocytes but not in neurons. **a.** Experimental design: two groups of 2-months old mice were used in the study. The control group (Ctrl) received standard chow; the test group received HFD for one month. **b.** Weight change in Ctrl and HFD. **c.** Percentage of body fat in Ctrl and HFD mice after one month of diet. **d.** An example of Raman spectrum from astrocytic soma showing the peaks corresponding to different molecules and states of the molecules. Identified peaks show the Raman shift values and the name of the corresponding molecule. Cyts *c,b*(Fe^2+^) – reduced cytochromes *c* and *b*, Phe – phenylalanine. **e-j**, Relative amounts of lipids (e), cholesterol (f), and reduced cytochromes *c* and *b* (g) in astrocytes and (h, I, j – correspondingly) in neurons in control and after HFD. The data presented as mean ± SEM; circles are individual measurements for different mice in control (empty) and HFD (orange) groups; N.S. p > 0.05, *p < 0.05, **p <0.01, ***p < 0.001, two-sample *t*-test.

Fatty acids serve both as an energy source and building blocks for lipids. Hence, we tested the effect of HFD on the metabolic status of astrocytes and neurons in the hippocampus with Raman microspectroscopy. This technique quantifies vibrational modes of molecules, thus revealing their relative content (e.g., lipids or proteins) and state (e.g., redox; Fig. 1d and S2). Astrocytes and neurons were identified with double immunocytochemical staining of hippocampal slices. Exposure to HFD significantly increased the relative amount of lipids in astrocytes (lipids/proteins ratio in control: 0.74 ± 0.03, n = 6; in HFD: 1.22 ± 0.04, n = 9; p < 0.001, two-sample t-test; Fig. 1e). Notably, there was a trend (although the difference did not reach significance) for an increase in cholesterol content (cholesterol/phosphatidylcholine ratio in control: 1.03 ± 0.04, n = 5; in HFD: 1.27 ± 0.12, n = 7; p = 0.14, two-sample t-test; Fig. 1f). The relative amount of reduced *c*- and *b*-type cytochromes was significantly higher in astrocytes after HFD (Cyts *c,b(Fe^2+^)*/phenylalanine ratio in control: 2.6 ± 0.1, n = 6; in HFD: 3.2 ± 0.2, n = 9; p = 0.037, two-sample t-test; Fig. 1g, S2). Detailed analysis of characteristic selective Raman peaks of *c*- and *b*-type cytochromes in astrocytes demonstrate that the overall increase in the relative amount of their reduced forms is due to the significant increase in the relative content of reduced c-type cytochromes (Fig. S2). In contrast, HFD significantly affected neither the relative amount of lipids and cholesterol nor the amount of reduced *c,b*-type cytochromes in neurons (Fig. 1h-j, S2). Therefore, HFD triggers metabolic changes in hippocampal astrocytes but not in neurons. This diet increased the synthesis of lipids serving as building blocks for astrocytic membrane. The observed increase in the relative amount of reduced *c*- and *b*-type cytochromes in astrocytes of HFD mice denotes the overloading of electron transport chain (ETC) with electrons, presumably be due to the increased amount of primary electron donors coming from β-oxidation of fatty acids. The rise in the amount of reduced electron carriers in the respiratory chain may increase the generation of reactive oxygen species (ROS). Astrocytic ROS have been recently implicated in the modulation of brain metabolism and mouse behavior (Vicente-Gutierrez et al., 2019).

### 2. HFD promotes astrocytic enlargement

Enhanced membrane biogenesis may promote astrocyte growth following HFD. To test this hypothesis, we loaded CA1 *str. radiatum* astrocytes in hippocampal slices with a fluorescent dye, Alexa Fluor 594, through a patch pipette (Popov et al., 2021). The three-dimensional (3D) reconstruction of stained astrocytes was made from z-stacks of images obtained with two-photon microscopy (Fig. 2a). Astrocytic branches were traced, and 3D Sholl analysis was performed with homemade Python code (Fig 2b. and see Method section). The number of primary branches emanating from soma was not significantly affected by HFD (control: 7.16 ± 0.60, n = 6; HFD: 8.14 ± 0.55, n = 7; p = 0.12, two-sample t-test; Fig. 2c), but the maximal number of intersections (control: 20.8 ± 1.5, n = 6; HFD: 25.1 ± 1.5, n = 7; p = 0.03, two-sample t-test; Fig. 2d) and mean branch length (control: 6.3 ± 0.3 μm, n = 6; HFD: 7.7 ± 0.2 μm, n = 7; p = 0.002, two-sample t-test; Fig. 2e) were increased. These changes enlarged astrocytic territorial domain area measured as a projection of imaged astrocytes along the z-axis (control: 1998 ± 257 μm^2^, n = 6; HFD: 2723 ± 271 μm^2^, n = 7; p = 0.04, two-sample t-test; Fig. 2f,g).

**Figure 2.**
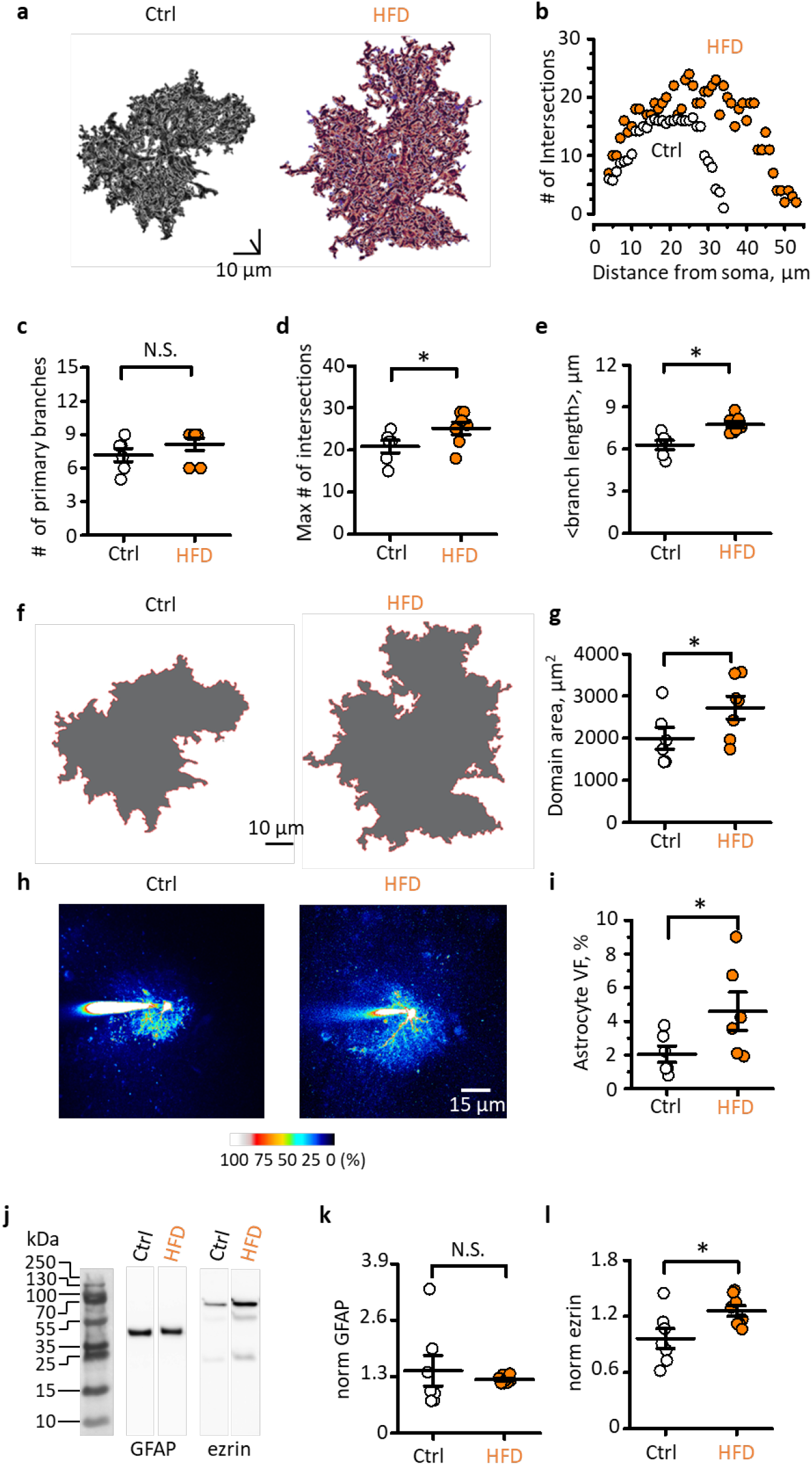
HFD promotes astrocyte growth and enlarges the astrocytic domain. **a.** Three-dimensional (3D) reconstruction of control (Ctrl) and HFD astrocytes loaded with fluorescent dye via patch pipette. **b.** The number of intersections of astrocytic branches with concentric spheres (3D Sholl analysis) for astrocytes presented in **a**. **c.** The number of primary branches of astrocytes in control and after HFD. **d.** The maximal number of intersections in control and after HFD. **e.** Mean branch length in control and after HFD. **f.** Astrocytic territorial domains obtained as a projection of astrocytes along the z-axis. **g.** Areas of the astrocytic territorial domains in control and after HFD. **h.** Astrocytes loaded with Alexa Fluor 594 through patch pipette in control and after HFD. The fluorescence intensity is normalized to the fluorescence of soma to illustrate astrocytic VF distribution. **i.** Astrocytic VF in control and after HFD. **j.** Representative Western blots of the mouse hippocampus homogenates stained by antibodies against GFAP and ezrin. **k,l** GFAP (k) and ezrin (l) protein level normalized to total protein level in control and after HFD. The data presented as mean ± SEM; circles are individual measurements for different mice in control (empty) and HFD (orange) groups; N.S. p > 0.05, *p < 0.05, two-sample *t*-test.

Two-photon imaging allows visualization of astrocytic branches and territorial domain, while leaflets remain beyond resolution (Semyanov and Verkhratsky, 2021). Hence, we analyzed volume fraction (VF) occupied by astrocytic leaflets as a ratio of fluorescence of unresolved processes to the fluorescence of soma (Fig. 2h and S3) (Medvedev et al., 2014; Minge et al., 2021; Plata et al., 2018). The VF of leaflets has significantly increased following HFD (control: 2.0 ± 0.5 %, n = 6; HFD: 4.6 ± 0.1 %, n = 6; p = 0.03, two-sample t-test; Fig. 2i).

Astrocytic enlargement is often considered to be a sign of reactive astrogliosis associated with increased expression of the glial fibrillary acidic protein (GFAP) (Escartin et al., 2021). However, no significant change in GFAP protein was observed in hippocampal tissue following HFD (control: 1.4 ± 0.4, n = 7; HFD: 1.23 ± 0.03, n = 8; p = 0.3, two-sample t-test; Fig. 2j,k, and S4). In contrast, Western blotting revealed an increased expression of ezrin implicated in astrocytic growth (control: 0.96 ± 0.11, n = 7; HFD: 1.26 ± 0.06, n = 8; p = 0.013, two-sample t-test; Fig. 2j,l, and S4).

Astrocytic enlargement induced by HFD was not associated with a change in the number of astrocytes stained with sulforhodamine 101 (Fig. S5). Thus, individual astrocytes increased their presence in the brain active milieu through extended space coverage by branches and leaflets. The expansion of astrocytic processes could affect the coupling of these cells through gap junctions. We counted the number of coupled astrocytes labeled by Alexa Fluor 594 diffusing through gap junctions from an individual patch-pipette loaded astrocyte to test this possibility. No significant difference was observed in the number of connected cells between control and HFD groups (Fig. S5).

### 3. HFD enhances glutamate uptake by astrocytes

Astrocyte enlargement is associated with an expansion of the cell membrane surface, increasing membrane capacitance, and total conductance (if channel density is unchanged). The slope of current-voltage relationships from current (ΔI) generated in response to voltage steps (ΔV) in CA1 *str. radiatum* astrocytes control and HFD mice (Fig. 3a,b) was used to determine cell input resistance (Ri reciprocal of membrane conductance). Consistent with larger membrane surface, Ri tended to be smaller in astrocytes after HDF, but the difference with control did not reach significance (control: 25 ± 5 MΩ, n = 6; HFD: 17 ± 3 MΩ, n = 10; p = 0.14, two-sample t-test; Fig. 3c).

**Figure 3.**
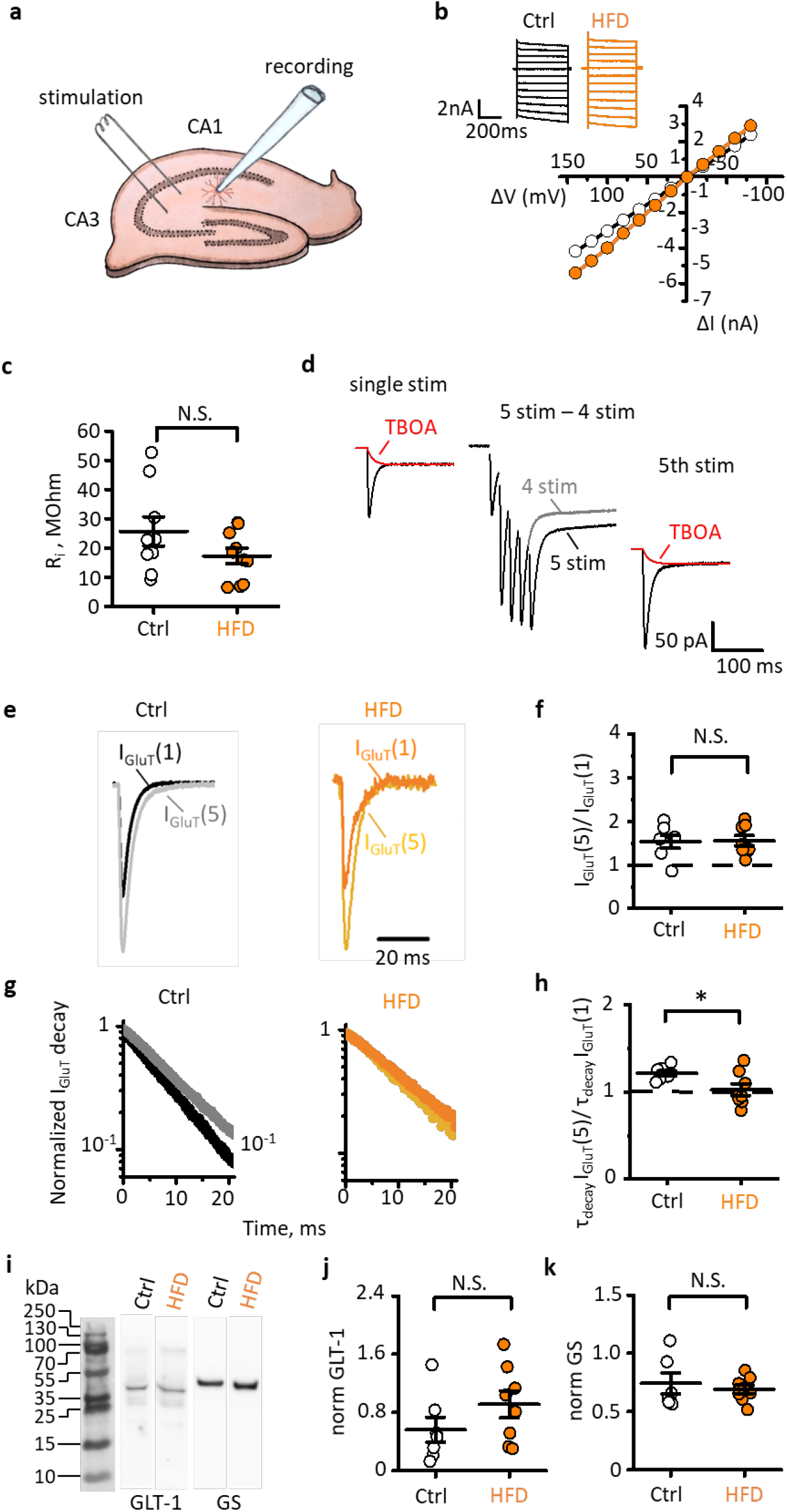
HFD reduces glutamate spillover. **a.** Experimental design: recording of currents in voltage-clamped CA1 *str. radiatum* astrocytes. Synaptically induced currents were triggered with the extracellular stimulating electrode. **b.** Current-voltage relationship for representative astrocytes in control (Ctrl) and after HFD. *Inset*, currents recorded in response to voltage steps. **c.** Input resistance (Ri) of astrocytes in control and after HFD. **d.** Astrocytic currents were recorded in response to a single stimulus, five and four (grey trace) stimuli (50 Hz) delivered via an extracellular electrode placed on Schaffer collaterals. Response to four stimuli was subtracted from response to five stimuli to isolate the response to the fifth stimulus. DL-TBOA was added at the end of the experiment, and residual current (red trace) was tail-fitted and subtracted from the astrocytic response to isolate the transporter current (I_GluT_). **e.** Superimposed I_GluT_(1) and I_GluT_(5) in control and after HFD. **f.** I_GluT_(5)/I_GluT_(1) ratio in control and after HFD. **g.** The normalized decay phase of I_GluT_(1) (darker color) and I_GluT_(5) (lighter color) in control and after HFD in semilogarithmic scale. **h.** τ_decay_ I_GluT_(5)/τ_decay_ I_GluT_(1) ratio in control and after HFD. **i.** Representative Western blots of the mouse hippocampus homogenates stained by antibodies against GLT-1 and GS. **j,k.** GLT-1(j) and GS (k) protein level normalized to total protein level in control and after HFD. The data presented as mean ± SEM; circles are individual measurements for different mice in control (empty) and HFD (orange) groups; N.S. p > 0.05, *p < 0.05, two-sample *t*-test.

Increased VF of leaflets arguably translates into enhanced astrocytic coverage of synapses, leading to a more efficient uptake of synaptically released glutamate. To test this possibility, we recorded synaptically induced currents in CA1 *str. radiatum* astrocytes in response to stimulation of Schaffer collaterals with a single stimulus or with four and five stimuli at 50 Hz (Fig. 3a,d). Isolated response to the fifth stimulus was obtained by subtracting the response to four stimuli from response to five stimuli. The experiments were performed in the presence of AMPA (α-amino-3-hydroxy-5-methyl-4-isoxazolepropionic acid) and NMDA (N-methyl D-aspartate) receptor blockers to prevent postsynaptic K^+^ efflux (Shih et al., 2013). At the end of each experiment, glutamate transporter current was blocked with a selective inhibitor DL-TBOA (DL-threo-β-hydroxyaspartic acid). The residual current was tail-fitted and subtracted from synaptic currents to isolate net transporter currents. Activity-dependent facilitation of glutamate release was measured as a ratio between transporter currents to the fifth stimulus and a single stimulus (I_GluT_(5)/I_GluT_(1), Fig. 3e). The ratio did not differ significantly between astrocytes of control and HFD mice (control: 1.5 ± 0.1, n = 7; HFD: 1.6 ± 0.1, n = 8; p = 0.45, two-sample t-test; Fig. 3f). This result suggests that synaptic release probability was not affected by HFD.

Next, we analyzed the decay time (τ_decay_) of transporter current, which correlates with the timecourse of glutamate clearance (Diamond, 2005). The decays of both I_GluT_(1) and I_GluT_(5) were linear in semilogarithmic scale (Fig. 3g) and therefore were fit with monoexponential functions. Activity-dependent spillover of glutamate was quantified as τ_decay_ I_GluT_(5)/ τ_decay_ I_GluT_(1) ratio. This ratio was significantly reduced after HFD (control: 1.21 ± 0.03, n = 7; HFD: 1.03 ± 0.07, n = 8; p = 0.015, two-sample t-test; Fig. 3h), suggesting that increased astrocytic presence in synaptic microenvironment reduces glutamate spillover in the hippocampus.

Glutamate spillover can be potentially reduced because of increased expression of glutamate transporters or glutamine synthetase (GS), which converts glutamate to glutamine. Analysis of protein expression did not reveal significant changes in the expression of glutamate transporter-1 (GLT-1; control: 0.6 ± 0.2, n = 7; HFD: 0.9 ± 0.2, n = 8; p = 0.09, two-sample t-test; Fig. 3i,j, and S4) and GS (control: 0.74 ± 0.09, n = 6; HFD: 0.69 ± 0.04, n = 8; p = 0.28, two-sample t-test; Fig. 3i,k, and S4) in the hippocampus of HFD mice ruling out such possibility. However, we observed a trend towards a higher expression level of GLT-1, which may reflect a need for more transporters on a larger membrane surface after HFD.

### 4. HFD enhances hippocampal LTP and promotes exploratory behavior

When glutamate spillover is reduced, fewer extrasynaptic NMDA receptors are activated, thus enhancing long-term potentiation (LTP) (Popov et al., 2020; Valtcheva and Venance, 2019). Field excitatory postsynaptic potentials (fEPSPs) were recorded with extracellular glass electrode placed in CA1 *str. radiatum* of hippocampal slices. The LTP, measured as an enhancement of fEPSP amplitude, was triggered by high-frequency stimulation (HFS) of Schaffer collaterals with an extracellular electrode. The LTP magnitude was significantly higher in slices obtained from HFD mice than in control (control: 142 ± 5 % of baseline, n = 7; HFD: 162 ± 10 % of baseline, n = 6; p = 0.044, two-sample t-test; Fig. 4a,b). To test if LTP enhancement was due to more efficient glutamate uptake, we partially blocked glutamate transporters with 100 nM TFB-TBOA. In the presence of TBOA the magnitude of LTP in both control and HFD mice was indistinguishable between two groups (control: 130 ± 5 % of baseline, n = 8; HFD: 132 ± 5 % of baseline, n = 10; p = 0.36, two-sample t-test; 4c,d). This suggests that enhanced uptake of synaptically released glutamate promotes hippocampal LTP in HFD mice.

**Figure 4.**
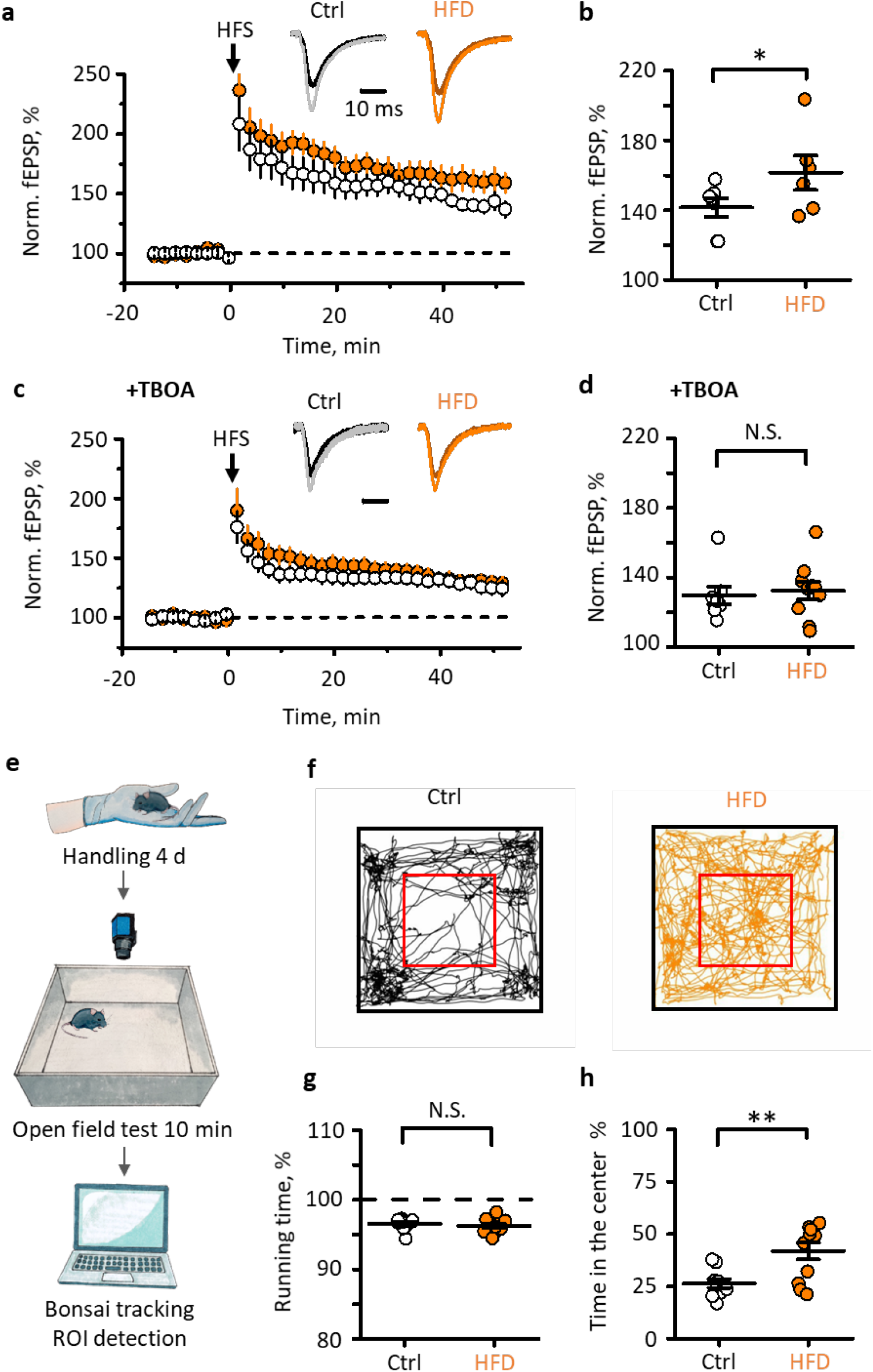
HFD enhances LTP and promotes exploratory behavior. **a.** LTP in CA1 *str. radiatum* in response to high-frequency stimulation (HFS) of Schaffer collaterals. The timecourses of fEPSP amplitudes in control and HFD slices are normalized to baseline before HFS. *Inset*, fEPSPs recorded at baseline (dark colors) and one hour after HFS (light colors). **b.** The magnitude of LTP one hour after HFS. **c.** Same as in **a**, but in the presence of TFB-TBOA. **d.** Same as in **b**, but in the presence of TFB-TBOA. **e.** Mice were handled for four days before they were placed in a square open field test. In the test, the animals were left for ten minutes, and their movements were recorded with the video camera for subsequent tracing with open-source software Bansai **(Lopes and Monteiro, 2021)**. **f.** The tracks obtained for control and HFD mice. The red box defines the center. **g.** Percentage of time the mouse was running for control and HFD groups. **h.** Percentage of time the mouse spent in the center of the arena for control and HFD groups. The data presented as mean ± SEM; circles are individual measurements for different mice in control (empty) and HFD (orange) groups; N.S. p > 0.05, *p < 0.05, **p < 0.01, two-sample *t*-test.

Enhanced glutamate uptake and synaptic plasticity associated with astrocytic hypertrophy could affect animal behavior. To test this possibility, we performed an open field test assaying gross locomotor activity, anxiety, and exploratory behavior (Fig. 4e,g). No significant difference in locomotor activity measured as percentage of active time was detected between two groups of animals (control: 96 ± 3 %, n = 10; HFD: 96 ± 3 %, n = 11; p = 0.48, two-sample t-test; 4f). However, HFD animals spent significantly more time in the center of the arena (control: 26 ± 2 %, n = 10; HFD: 42 ± 4 %, n = 11; p = 0.002, two-sample t-test; 4g), indicating a lower level of anxiety or higher willingness to explore or both.

## Discussion

We found that one month of high-fat nourishment of young mice changes the metabolism of hippocampal protoplasmic astrocytes but not neurons. This increases astrocytic morphological complexity and enhances glutamate clearance. Mice exposed to this nourishment regime increased their weight without becoming obese. Raman microspectroscopy with single-cell resolution revealed increased lipid content and relative amount of reduced cytochromes in astrocytes and not in neurons following HFD, indicating that astrocytes are the primary target of nutritionally delivered fatty acids in the brain. Protoplasmic astrocytes in the hippocampus of fat-nourished mice became larger and more complex, resulting in the expansion of VF occupied by leaflets interacting with synapses (Semyanov and Verkhratsky, 2021). The increase in astrocytic synaptic coverage promoted glutamate clearance, reduced glutamate spillover, and enhanced LTP. These cellular changes paralleled the elevated exploratory activity of HFD animals.

Enlargement of astrocytes following HFD is consistent with a recent report showing that an obesogenic ‘cafeteria diet’ that also contains high amounts of fat triggers astrocytic hypertrophy (Lau et al., 2021). However, increased astrocytic presence in the brain active milieu observed in our experiments is distinct from hypertrophy associated with reactive astrogliosis, which by definition is a response to pathological insults (Escartin et al., 2021). Astrocyte reactivity is commonly associated with increased expression of GFAP, which was not observed following HFD. Instead, our experiments revealed upregulation of ezrin. This protein binds the actin cytoskeleton to the plasma membrane playing a role in astrocyte growth (Zhou et al., 2019). Physiological enlargement influences astrocytic homeostatic capabilities. For example, enlargement of astrocytes following caloric restriction enhances K^+^ clearance and glutamate uptake leading to elevated LTP (Popov et al., 2020).

Conversely, aging-related shrinkage of astrocytes impairs glutamate uptake, which consequently attenuates LTP (Popov et al., 2021). Thus, the cumulative effect of environmental factors (generally referred to as an exposome) affects astrocytes morphology affecting branches, leaflets, and domain size, consequently modifying other elements of the brain active milieu. Similarly, HDF links astrocytic metabolism and morphological remodeling to cellular function and behavioral output.

Exposome defines the brain metabolism by targeting astrocytes in particular. Physical exercise, as well as exposure to an enriched environment, potentiate astrocytic growth, which increases synaptic coverage and homeostatic support, ultimately modifying learning, memory, and behavior (Augusto-Oliveira and Verkhratsky, 2021; Rodriguez et al., 2013). Likewise, dieting changes astrocytic morphology and function ((Popov et al., 2020) and the present study). The cognitive and behavioral outcomes of lifestyle seem to be age-dependent, with similar diets triggering either beneficial or adverse effects in different ages. The developing brain has an excessive metabolic rate consuming 50% of the body’s energy (Bonvento and Bolanos, 2021), with fatty acids metabolism contributing to energy supply (Steiner, 2019). Excessive dietary fat appears an essential component of the diet for children and young adults receiving training and education. At advanced ages, the HFD may trigger numerous systemic pathologies, affecting the brain and limiting cognitive longevity (Pistell et al., 2010). Astrocytes, being a homeostatic center of the active milieu of the brain, translate diets to brain circuitry and behavior.

## Methods

### Animals

All procedures were performed in accordance with Rus-LASA ethical recommendations approved by the Institute of Bioorganic Chemistry ethical committee. The 2-month-old C57BL/6 male mice were divided into control and HFD groups. The HFD group received the fat-enriched food (SNIFF, Germany, see Table S1) and the control animals had the standard chow for 1 month prior to the experiments. Both groups had access to food and water *ad libitum*. For behavioral testing, all animals were held in groups on a reversed light-dark cycle (light was off at 10 AM, on at 10PM).

After behavioral testing, some mice from control (n = 5) and the HFD (n = 7) groups were sacrificed by cervical dislocation for further autopsy. Subcutaneous and abdominal fat tissue and visceral organs were isolated and weighed with electronic scales Scout Pro SP202 (Ohaus, Switzerland). Fat percentage was calculated as a ratio of subcutaneous and abdominal fat tissue weight to the bodyweight of the mouse.

### Hippocampal slices

The mice were anesthetized with (1-chloro-2,2,2-trifluoroethyl-difluoromethyl ether) before being sacrificed. The brain was removed and immersed in an ice-cold solution containing (in mM): 50 sucrose, 87 NaCl, 2.5 KCl, 8.48 MgSO_4_, 1.24 NaH_2_PO_4_, 26.2 NaHCO_3_, 0.5 CaCl_2_, and 22 D-glucose. Then hippocampus was removed and cut into 350 μm thick slices with a vibrating microtome (Microm HM650V, Thermo Scientific, USA). Slices were left to recover for 1 hour at the temperature of 34°C in a recovery solution containing (in mM): 119 NaCl, 2.5 KCl, 1.3 MgSO_4_, 1 NaH_2_PO_4_, 26.2 NaHC O_3_, 1 CaCl_2_, 1.6 MgCl_2_, 22 D-glucose. For electrophysiology and two-photon fluorescence imaging experiments, the slices were transferred into an immersion chamber where were continuously perfused (2 ml/min) at the temperature of 34°C with artificial cerebrospinal fluid containing (in mM): 119 NaCl, 2.5 KCl, 1.3 MgSO_4_, 1 NaH_2_PO_4_, 26.2 NaHCO_3_, 2 CaCl_2_, and 11 D-glucose. All solutions were saturated with carbogen: 95% O_2_ and 5% CO_2_, and had an osmolarity of 295 ± 5 mOsm and pH of 7.4. For result consistency and to reduce the number of animals used in this study, hippocampal slices from the same animal were used for Raman spectroscopy, electrophysiology, and morphology analysis.

### Immunocytochemical staining of hippocampal slices for Raman microspectroscopy

Immunocytochemical staining of brain slices for GFAP was used to identify astrocytes for the selective study of mitochondria redox state, proteins, and lipids amount in astrocyte cytoplasm. The staining was performed in following steps: (1) acute hippocampal slices after the recovery period were placed into 4% paraformaldehyde (PFA) solution (37°C) for 60 min and then washed twice in phosphate buffer solution (PBS); (2) PFA-fixed slices were transferred into PBS + 0.3% Triton-X100 solution for 20 min and then into PBS + 0.1% Tween 20 + 5% BSA solution (25°C) for 90 min; (3) the slices were then incubated in the primary antibody solution (anti-glial fibrillar acidic protein (GFAP) rabbit polyclonal antibody, Abcam, catalog number ab7260; and anti-neuronal nuclear protein (NeuN) chicken polyclonal antibody, Novusbio catalog number NBP2-104491) in PBS + 0.01% Tween 20 for 60 h at 25°C; (4) slices were washed trice in PBS, 10 min, 25°C; (5) slices were incubated in the secondary antibody solution (Alexa Fluor 488 AffiniPure donkey anti-rabbit IgG (H+L), Jackson ImmunoResearch, catalogue number 711-545-152 and Cy-5 AffiniPure donkey anti-chicken IgG (H+L), Jackson ImmunoResearch, catalog number 703-175-155) for 2 h at 25°C; (5) slices were washed twice in PBS and in trice in deionized water. Then the slices were placed on the glass slide and dried in the dark. Stained slices were stored at room temperature in the dark. The secondary antibodies were chosen on the basis of their spectral properties (excitation and emission wavelengths, Alexa Fluor 488: λ_ex_=490 nm; λ_em_=510 nm and Cy-5: λ_ex_=650 nm; λ_em_=670 nm) so they did not emit fluorescence with the 532 nm laser excitation, which was used for Raman spectroscopy.

### Analysis of mitochondria redox state and relative amount of lipids in astrocyte cytoplasm with Raman microspectroscopy

Confocal Raman microspectrometer NTEGRA SPECTRA (NT-MDT, Zelenograd, Russia) with the inverted Olympus microscope was used to record Raman spectra of GFAP-stained astrocytes and NeuN-stained neurons in the hippocampus of mouse brain slices. Astrocytes and neurons were identified by the Alexa Fluor 488 and Cy-5 fluorescence, respectively, in the epifluorescence mode of the Olympus microscope with the halogen lamp illumination. After the localization of either astrocyte or neuron, the Raman microspectrometer was switched to the Raman mode and the Raman spectrum was recorded from the identified cell using laser excitation of 532 nm with the x40 NA0.45 objective. This excitation wavelength produced negligible fluorescence of Alexa Fluor 488 or Cy-5, which did not interfere with Raman scattering of cells. The laser power per registration spot was no more than one mW and the spectrum accumulation time was 60 s. Raman spectra were recorded from the cell soma. The diameter of the registration spot in the lateral plane was appr. 700 nm, and the height of the registration volume was appr. 1.5-2 μm.

### Analysis of Raman spectra

Raman spectra were analyzed with open-source software Pyraman, available at https://github.com/abrazhe/pyraman. The baseline was subtracted in each spectrum. It was defined as a cubic spline interpolation of a set of knots, number and x-coordinates of which were selected manually outside any informative peaks in the spectra. The number and x-coordinates of the knots were fixed for all spectra in the study. y-coordinates of the knots were defined separately for each spectrum as 5-point neighborhood averages of spectrum intensities around the user-specified x-position of the knot. The parameters for baseline subtraction were chosen after the processing of approximately 40 spectra from different cells. After the baseline subtraction, the intensities of peaks with the following maximum positions were defined: 700, 720, 750, 1003, 1445, and 1660 cm^-1^. These peaks correspond to vibrations of bonds in cholesterol, phosphatidylcholine, and reduced cytochromes of *b*- and *c*-types, Phe-residues, lipids, and proteins, respectively (Brazhe et al., 2012; Harkness et al., 2012; Kakita et al., 2012; Love et al., 2020; Okada et al., 2012; Perry et al., 2017). Normalized Raman peak intensities were used to compare lipid amount in the cytoplasm and the redox state of mitochondrial cytochromes in control and HFD mice (Love et al., 2020; Perry et al., 2017):

- I_1445_/I_1660_ ratio corresponding to the relative amount of all lipids normalized on the total amount of proteins in the region of observation;
- I_700_/I_720_ ratio corresponding to the relative amount of cholesterol *vs*. amount of phosphatidylcholine;
- I_750_/I_1003_ ratio corresponded to the relative amount of both reduced *c*- and *b*-type cytochromes normalized on the total amount of phenylalanine;
- I_604_/I_1003_ and I_1338_/I_1003_ ratios demonstrating relative amounts of reduced *c*-type cytochromes and *b*-type cytochromes, respectively, normalized on the total amount of proteins.

### Astrocyte morphometry

Astrocytes were loaded with fluorescent dye Alexa Fluor 594 through the patch pipette. Fluorescence was excited in two-photon mode, and images were collected with the Zeiss LSM 7 MP microscope equipped with Ti:sapphire femtosecond laser Chameleon Vision II.

### 3D Sholl analysis

z-stacks of images were first corrected for a constant fluorescence offset value and resampled to a uniform scaling along all axes by a linear interpolation along the z-axis. Offset subtraction was required for volume fraction (VF) analysis; offset value was estimated by a bootstrapping procedure as the x-axis intercept of the linear fit of fluorescence intensity variance vs. mean intensity in a large number of small randomly chosen blocks of the z-stack. Specifically, 10^5^ 10×10×10 blocks were chosen randomly from the z-stack as a representative sample of the data. Blocks with mean intensity higher than the top 95% were excluded from the further fit as they were likely to contain soma and primary branches of the astrocyte. Next, 100 sub-samples of 10% blocks were randomly chosen from the whole block collection and used for linear fits of intensity variance vs. mean intensity. The resulting slopes were treated as an empirical distribution of detector gain estimates and x-axis intercepts — as a distribution of offset value estimates. The mean of all the offset estimates resulted in the final offset value.

After offset subtraction and resampling, masks corresponding to astrocyte branches and processes were created. Elongated features corresponding to astrocyte processes were highlighted by using coherence-enhanced diffusion filtering (CEDF). CEDF was done sequentially in 2D planes along all three principal axes (i.e. XY, ZY, ZX planes), results for the three axes were averaged. Astrocyte processes were segmented by hysteresis thresholding with a low threshold set to one standard deviation (s.d.) of background noise and a high threshold set to three s.d. of background noise. The segmentation result was used for running the Sholl analysis plugin in ImageJ.

### Calculation of astrocytic VF

An image containing the middle of astrocyte soma was selected from the z-stack. The attention was paid that the fluorescence of soma was not saturated. Five cross-sections for each cell were plotted through the center of the soma at the angle of 72° from each other. Large fluctuations of fluorescence (> 10% of soma fluorescence and > 0.5 μm width) corresponding to astrocytic branches were cut out. VF was calculated as fluorescence intensity in unresolved processes normalized fluorescence intensity in the soma. Characteristic VF was obtained as mean VF at the distance of 20-25 μm from the soma.

### Western blotting

The hippocampi of control and HFD mice were frozen in liquid nitrogen. Then each hippocampus was separately homogenized in RIPA buffer containing SIGMAFAST protease inhibitor cocktail (Sigma-Aldrich, St. Louis, USA), diluted in loading buffer (120 mM Tris-HCl, 20 % [*v/v*] glycerol, 10 % [*v/v*] mercaptoethanol, 4 % [*w/v*] sodium dodecyl sulfate, and 0.05 % [*w/v*] bromophenol blue, pH 6.8), submitted to gel electrophoresis, and blotted onto nitrocellulose membranes (GE Healthcare, Chicago, USA). The membranes were blocked in 5% skim milk (Sigma-Aldrich) in TBS buffer (50 mM Tris, 150 mM NaCl, pH 7.4) + 0.2 % Tween-20 (Applichem, Darmstadt, Germany) for one h at room temperature and then incubated overnight at four °C with primary mouse antibodies for ezrin (Antibodies-Online, ABIN5542456, Aachen, Germany), with primary rabbit antibodies for GLT-1 (Abcam, 106289, Waltham, USA) or GFAP (Antibodies-Online, ABIN3044350), or with primary guinea pig antibodies for GS (Synaptic Systems, 367 005, Goettingen, Germany). After incubation with primary antibodies, membranes were rinsed in TBS with 0.2 % Tween-20 and incubated with HRP-conjugated secondary antibodies—anti-mouse IgG (Jackson Immunoresearch, 715-005-150, West Grove, USA), anti-rabbit IgG (Abcam, 6721), or anti-guinea pig IgG (Jackson Immunoresearch, 706-035-048) for one h. ECL substrate (Bio-Rad, Hercules, USA) was used for signal detection. Protein bands were visualized using an ImageQuant LAS500 chemidocumenter (GE Healthcare). The intensity of protein bands was quantified using the gel analyzer option of ImageJ software (NIH, Bethesda, USA). To exclude inter-sample variability, the averaged intensities of bands measured for each mice/protein were normalized to the average intensity of the lanes with total protein of the corresponding samples using the No-Stain labeling kit (A44449, Life Technologies, Carlsbad, USA).

### Electrophysiological recordings

Visually identified astrocytes were selected in the *str. radiatum* at 100 – 200 μm from the stimulating electrode and recorded in whole-cell mode with borosilicate pipettes (3 – 5 MΩ) filled with an internal solution containing (in mM): 135 KCH_3_SO_3_, 10 HEPES, 10 Na_2_phosphocreatine, 8 NaCl, 4 Na_2_-ATP, 0.4 Na-GTP (pH was adjusted to 7.2; osmolarity to 290 mOsm). 50 μM Alexa Fluor 594 (Invitrogen, USA) was added to the internal solution for morphological study. Passive astrocytes were identified by their small soma (5 – 10 μm in diameter), strongly negative resting membrane potential (around −80 mV), and linear current-voltage relationship. In the current-clamp mode, current steps were applied to corroborate the absence of membrane excitability. In voltage-clamp recordings, the astrocytes were held at −80 mV. Voltage steps of 20 mV (500 ms) from −140 mV to 80 mV were applied to obtain ΔI/ΔV relationships. Input resistance was determined by the slope of the ΔI/ΔV curve.

### Analysis of glutamate transporter current

One, four, and five electrical stimuli at 50 Hz were applied to Schaffer collaterals to trigger synaptic currents in the astrocytes, followed by a voltage step of −5 mV for monitoring cell input resistance. Signals were sampled at 5 kHz and filtered at 2 kHz. Current in response to the fifth stimulus was obtained by the subtraction of current evoked by four stimuli from current evoked by five stimuli.

The astrocytic currents induced by synaptic stimulation contain several components, including fast glutamate transporter current (I_GluT_) and slow K^+^ inward current (I_K_) (Sibille et al., 2014). I_GluT_ was obtained in the presence of antagonists of postsynaptic ionotropic receptors: 25 μM NBQX (2,3-Dioxo-6-nitro-1,2,3,4-tetrahydrobenzo[f]quinoxaline-7-sulfonamide disodium salt) – AMPA receptor antagonist, 50 μM D-APV (D-2-Amino-5-phosphonovaleric acid) – NMDA receptor antagonist, and 100 μM picrotoxin – GABA_A_ receptor channel blocker (all from Tocris Bioscience, UK). This cocktail prevented filed potential mediated component of astrocytic current and largely reduced I_K_ (Shih et al., 2013). After 10 minutes of recordings, 100 μM DL-TBOA (Tocris Bioscience, UK), an excitatory amino acid transporter (EAATs) blocker, was added to the bath to obtain residual IK, which was fit to the tail and subtracted from synaptic current to obtain pure I_GluT_. The I_GluT_ decay was fitted with a monoexponential function, and τdecay was calculated.

### Field potential recordings and LTP induction

The field excitatory postsynaptic potentials (fEPSPs) were recorded in CA1 *str.radiatum* with glass microelectrodes (resistance: 2–5 MOhm). For timecourse experiments, half-maximal stimulus intensity was chosen (the stimulus intensity when fEPSP amplitude was in 40–50% of the amplitude when the population spike appeared). The strength of stimulation was constant during the experiment, usually being 100–150 μA. The LTP was induced if the stable amplitude of the baseline fEPSP could be recorded for 15 min. Three trains of high-frequency stimulation (HFS, 20 pulses at 100 Hz, with an inter-train interval of 20 s protocol) were applied to induce LTP. The fEPSPs were recorded after induction protocol for at least 60 min. The LTP magnitude was estimated as the ratio of potentiated fEPSP amplitude (averaged in the interval of 50-60 min after the HFS) to baseline fEPSP amplitude (Popov et al., 2020).

### Open-field behavioral test

All behavioral testing was done between 10AM and 10PM during the animals’ active phase. Prior to the behavioral experiments, mice were handled for 4 days. On the test day, mice were habituated in the experimental room for 30 min prior to the start. All experiments were done in the darkroom with the red-light source invisible to mice and allowing for video recording with the GigE monochrome camera (The Imaging Source DMK 33GV024, Germany). Mice were placed in the center of a 40 × 40 × 40 cm open-field arena for a 10 min session and video recorded at a 50 fps rate. The arenas were cleaned between sessions with 70% ethanol, followed by distilled water. All video files were analyzed with a custom-made workflow in Bonsai (Open-source software, bonsai-rx.org). The workflow had two separate branches detecting the animal centroid (movement tracking) and the time spent in the arena center versus the periphery. The standard Bonsai nodes from OpenCV (computer vision) library were used to set the inverted binary threshold to detect a black mouse on the white background, followed by finding the largest binary region, a binary region analysis, and centroid node indicating the x and y position of the animal. We used the same threshold to estimate the time spent in the center of the arena, which was defined by the region of interest node, where white pixels were detected in each frame in the ROI activity detection node. To estimate the mice activity level, we used the speed threshold at 0.5 cm/s. The following statistical analysis of the output .csv files was performed with OriginPro (OriginLab Corp., USA).

### Statistics and reproducibility

All data are presented as the mean ± standard error of the mean (SEM). n – numbers indicate the number of animals. Statistical significance was assessed with Student’s t-test. p < 0.05 was considered statistically significant. Statistical analysis was performed by OriginPro (OriginLab Corp., USA).

## Acknowledgments

This work was supported by the Russian Foundation for Basic Research (RFBR), grant No. 21-54-53018.

## Supplementary information

**Figure S1.**
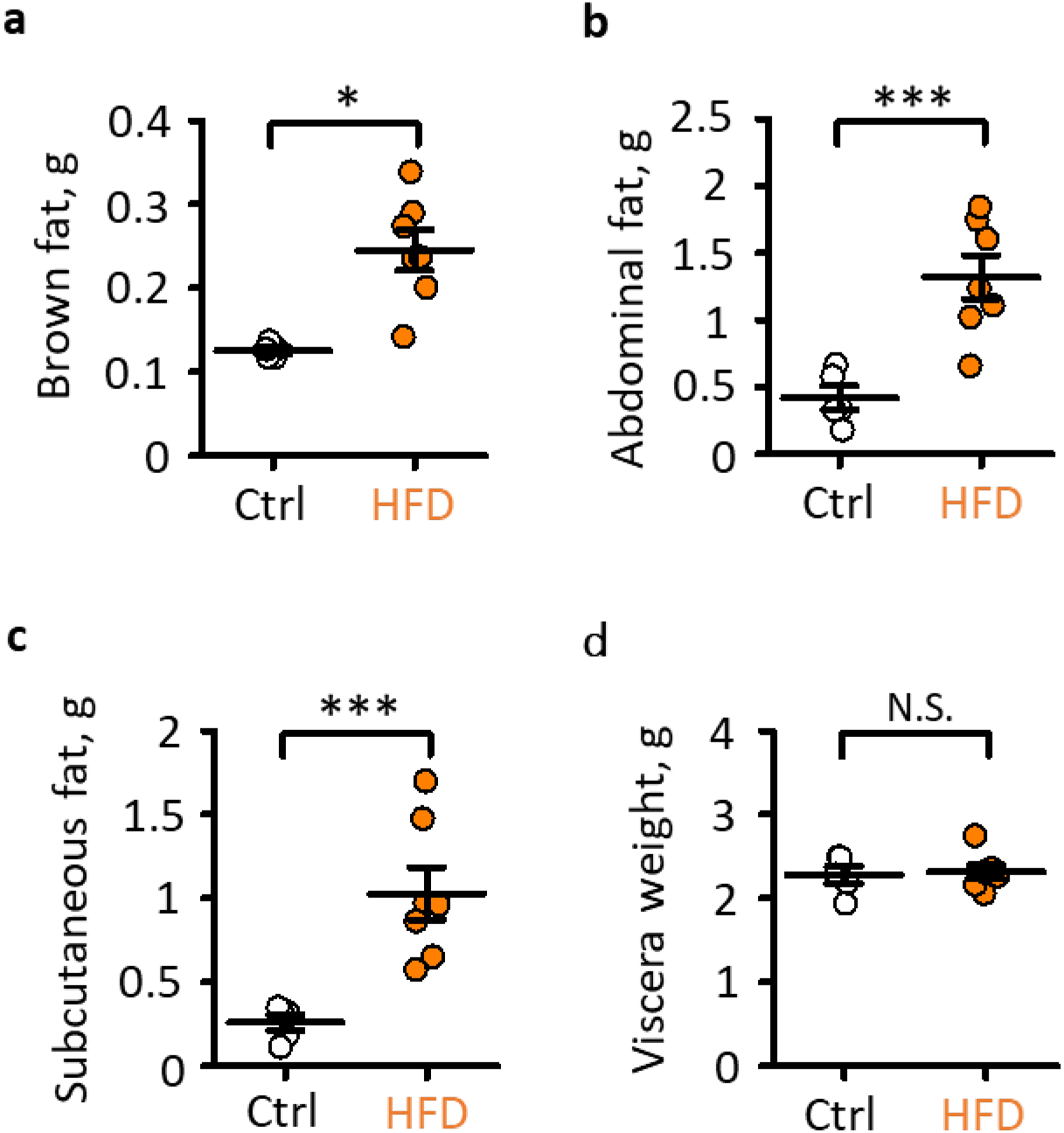
HFD increases all types of fat but not viscera weight. **a-c.** The weight of brown (a), abdominal (b), and subcutaneous (c) fat in control mice and mice after HFD. **d.** viscera weight in control mice and mice after HFD. The data presented as mean ± SEM; circles are individual measurements in control (empty) and HFD (orange) groups; N.S. p > 0.05, *p < 0.05, **p < 0.01, two-sample *t*-test.

**Figure S2.**
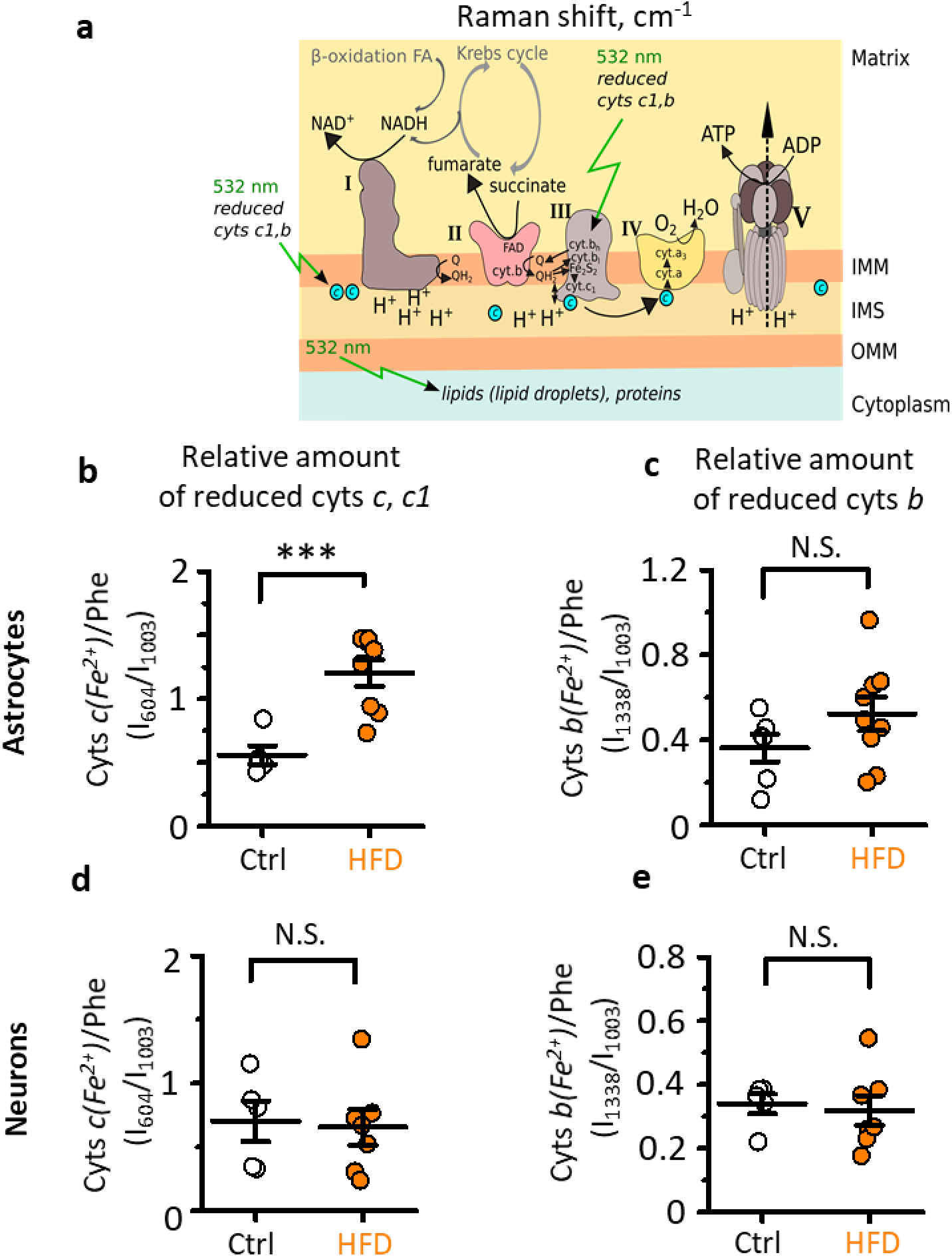
Changes in Raman spectral peaks related to reduced cytochromes c and b after HFD. **a.** Schematic illustrating the operation of electron transport chain highlighting the steps involving cytochromes *c* and *b*. **b,c** Relative amounts of reduced cytochromes *c, c1* (b), and *b* (c) in astrocytes in control and after HFD. **d,e.** Same as b,c but in neurons. The data presented as mean ± SEM; circles are individual measurements in control (empty) and HFD (orange) groups; N.S. p > 0.05, *p < 0.05, ***p < 0.001, two-sample *t*-test.

**Figure S3.**
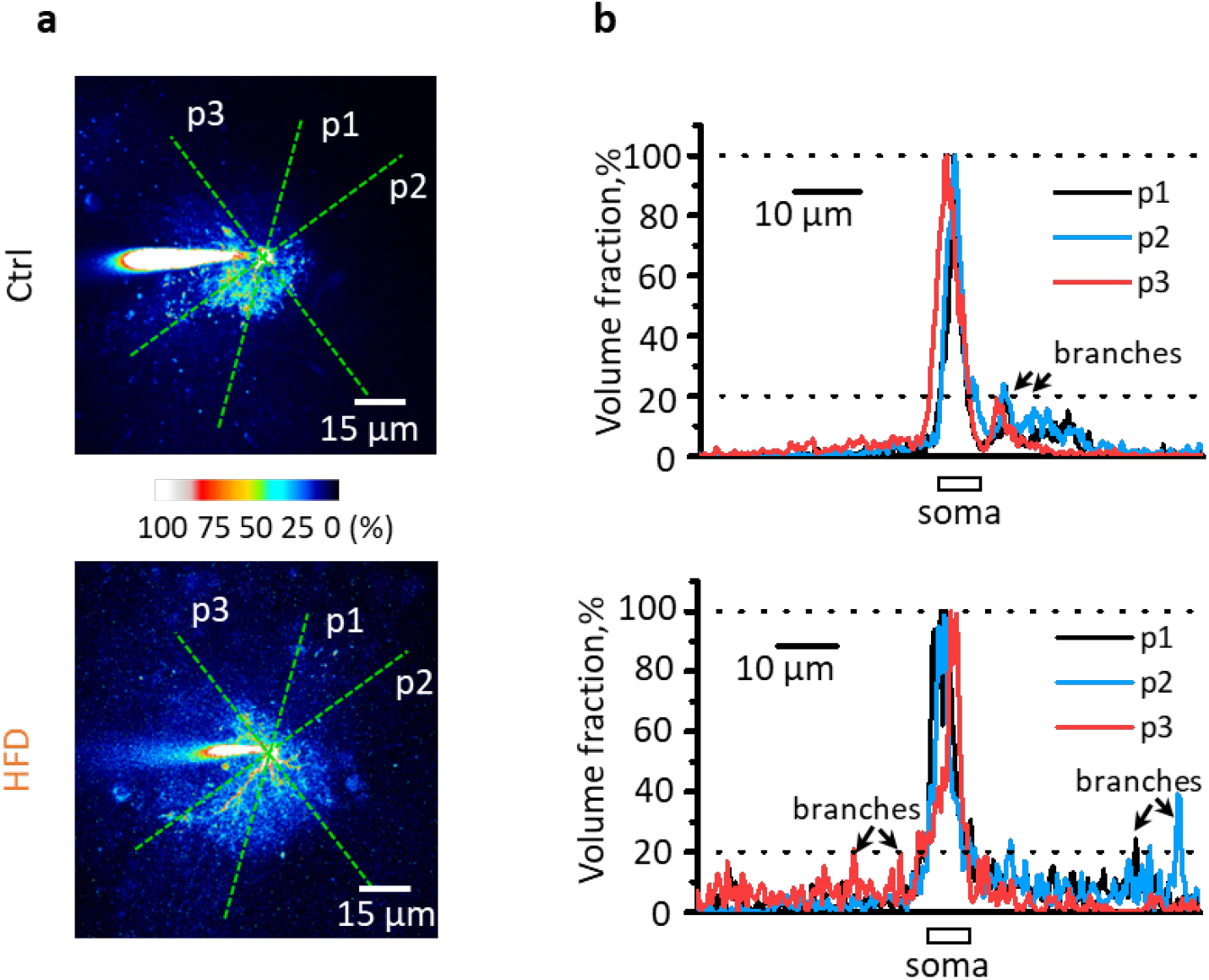
Measurements of astrocyte VF. **a.** Image fluorescence was normalized to fluorescence of astrocyte soma (100% of VF). Cross-sections through soma were made in different directions (p1-3). Top image – control, bottom – HFD. **b.** Fluorescence profiles were obtained for each cross-section. Fluorescence of soma and branches was excluded from the analysis. The remaining normalized fluorescence corresponds to VF of leaflets which was averaged for all cross-sections.

**Figure S4.**
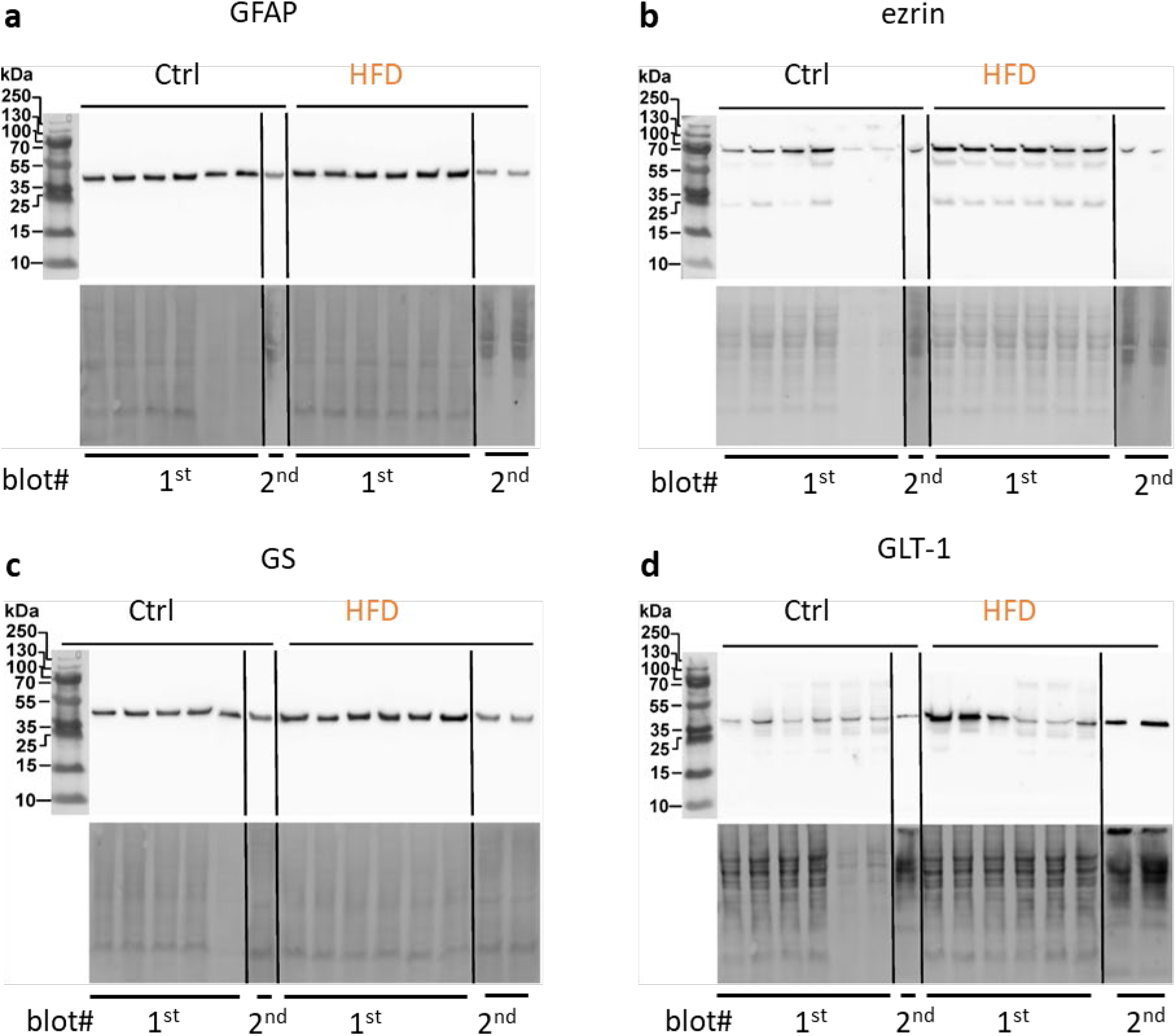
Western blot data for GFAP, ezrin, GS, and GLT-1. Individual blots for GFAP (a), ezrin (b) GS (c) and GLT-1 (d). The blots were done in two separate membranes as marked below (blot #). *Top*, the protein of interest. *Bottom*, total protein for normalization.

**Figure S5.**
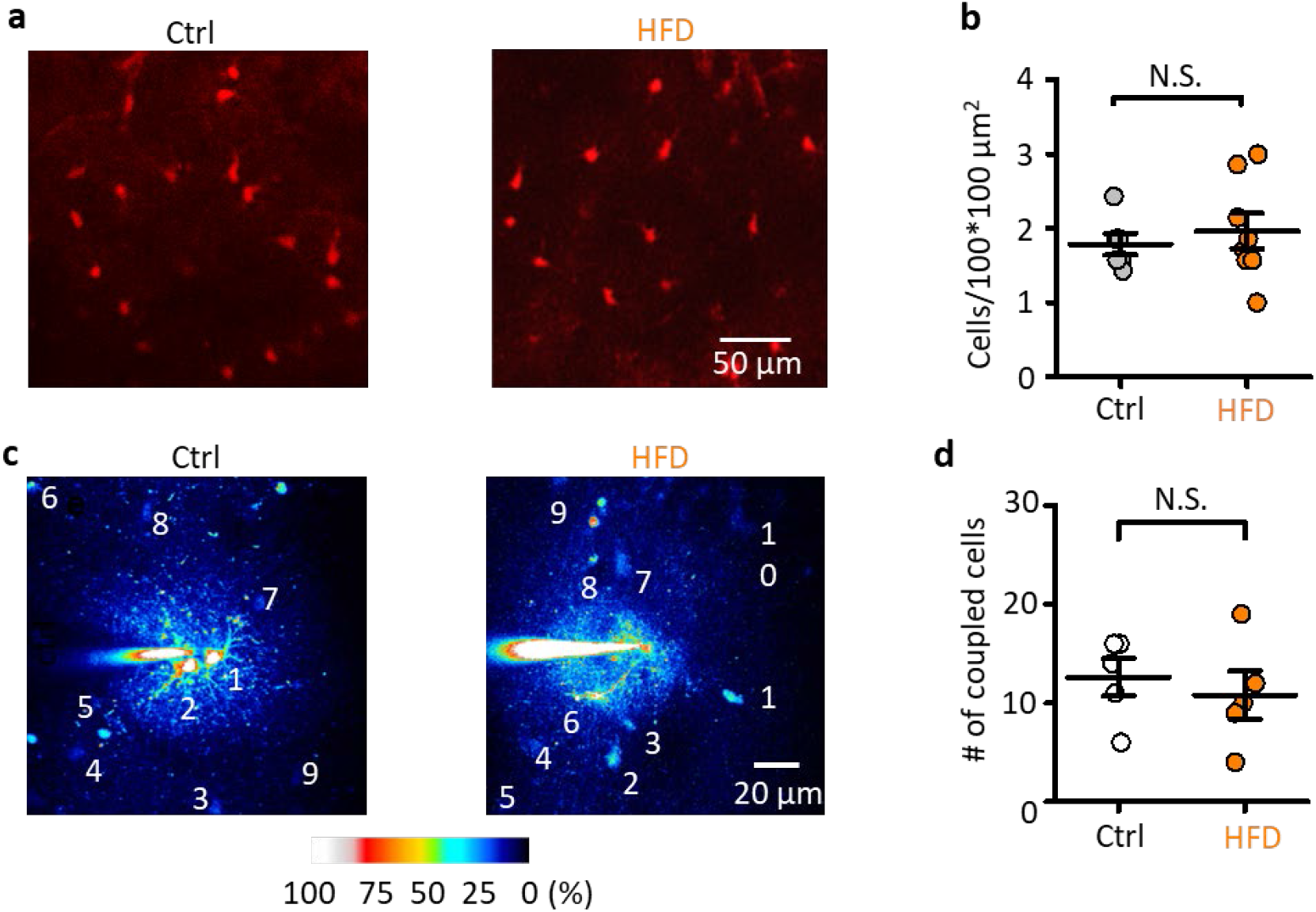
HFD does not affect astrocyte density or coupling to their neighbors. **a.** Astrocytes stained with sulforhodamine 101 (SR101) in control and after HFD. **b.** The density of astrocytes stained with SR101 in control and after HFD. **c.** Astrocytes loaded with Alexa Fluor 594 through patch pipette with neighboring astrocytes stained by the dye diffusion through gap junctions in control and after HFD. **d.** The number of coupled astrocytes stained with dye diffusion in control and after HFD. The data presented as mean ± SEM; circles are individual measurements in control (empty) and HFD (orange) groups; N.S. p > 0.05, two-sample t-test.

**Table S1.** Content of control chow and HFD.

| **Nutrients** | **Ctrl** | **HFD** |
| --- | --- | --- |
| Crude protein | 17.3% | 21% |
| Crude fat | 5.1% | 30.7% |
| Crude fibre | 5% | 5% |
| Crude ash | 3.9% | 5.4% |
| Strarch | 38.5% | 16.3% |
| Sugar | 11 % | 16.9% |
| Other | 19.2% | 4.7% |

